# The S-phase Cyclin Clb5 Promotes rDNA Stability by Maintaining Replication Initiation Efficiency in rDNA

**DOI:** 10.1101/2020.07.06.190892

**Authors:** Mayuko Goto, Mariko Sasaki, Takehiko Kobayashi

## Abstract

Regulation of replication origins is important for complete duplication of the genome, but the effect of origin activation on the cellular response to replication stress is poorly understood. The budding yeast ribosomal RNA gene (rDNA) forms tandem repeats and undergoes replication fork arrest at the replication fork barrier (RFB), inducing DNA double-strand breaks (DSBs) and genome instability accompanied by copy number alterations. Here we demonstrate that the S-phase cyclin Clb5 promotes rDNA stability. Absence of Clb5 led to reduced efficiency of replication initiation in rDNA but had little effect on the amount of replication forks arrested at the RFB, suggesting that arrival of the converging fork is delayed and forks are more stably arrested at the RFB. Deletion of *CLB5* affected neither DSB formation nor its repair at the RFB, but led to an accumulation of recombination intermediates. Therefore, arrested forks at the RFB may be subject to DSB-independent, recombination-dependent rDNA instability. The rDNA instability in *clb5*Δ was not completely suppressed by the absence of Fob1, which is responsible for fork arrest at the RFB. Thus, Clb5 establishes the proper interval for active replication origins and shortens the travel distance for DNA polymerases, which may reduce Fob1-independent DNA damage.

## INTRODUCTION

Precise duplication of the genome is crucial for maintaining genome integrity. However, DNA replication is constantly challenged by exogenous and endogenous stresses (1, 2). Obstacles such as DNA damage and tightly bound proteins cause the replication fork to stall. Stalled forks resume DNA replication when the replication stress is relieved or complete replication when a converging fork arrives (3). Failure to properly resolve a stalled fork results in chromosome rearrangements such as sequence deletion, duplication, or inversion, and chromosome translocations, which are hallmarks of cancer cells, cause human genomic disorders and influence genome diversity (2, 4).

The budding yeast ribosomal RNA (rDNA) region has a replication fork blocking site and undergoes genome rearrangements in response to replication fork stalling. Therefore, this region is useful for studying genome instability caused by replication-related recombination (5). The rDNA contains a tandem array of ∼150 copies of rDNA sequence at a single locus on chromosome XII (Fig. 1A). Each copy contains 35S and 5S rRNA transcription units, an origin of DNA replication, and a replication fork barrier (RFB) sequence. Although the origin in each copy has the potential to fire, DNA replication is initiated only from a subset of replication origins (6). DNA replication initially proceeds bi-directionally, but the replication fork moving against the 35S rDNA is stalled by Fob1 protein bound to the RFB site (7–9), potentially leading to DSB formation (10–12) (Fig. 1A). The rDNA copy number frequently changes in a manner mainly dependent on Fob1.

**Figure 1.**
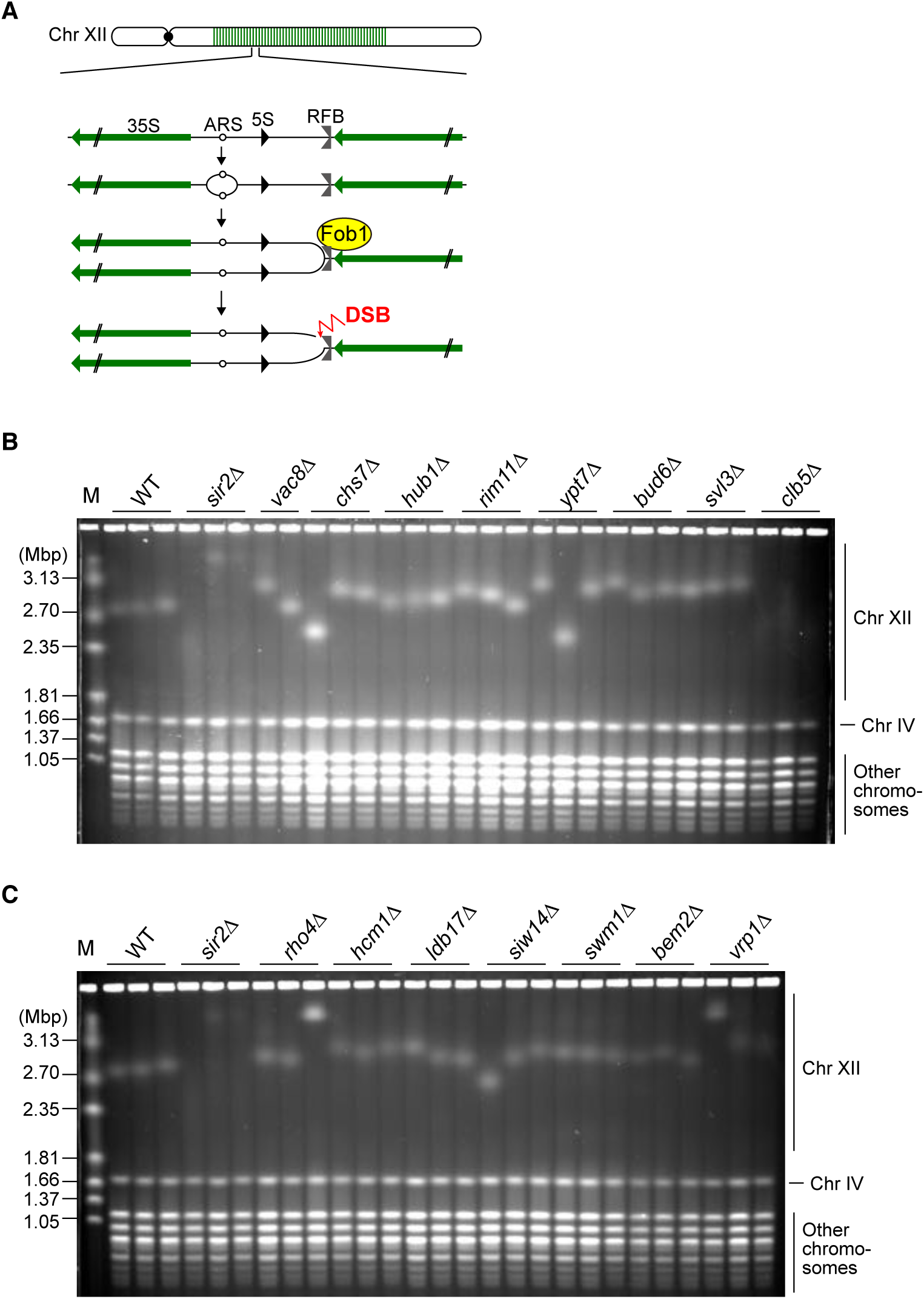
Absence of Clb5 causes rDNA instability. (A) DNA replication pattern in rDNA and changes in copy number. 35S, 35S rRNA; 5S, 5S rRNA; ARS, autonomously replicating sequence; RFB, replication fork barrier; DSB, DNA double-strand break. (B, C) PFGE analysis of the size heterogeneity of chromosome XII. The indicated genes were deleted from WT cells in the W303 background. DNA was extracted from three independent clones of WT and all mutant strains except for *vac8*Δ (two clones) and separated by PFGE. DNA was stained with ethidium bromide. M indicates *H. wingei* chromosomal DNA markers.

We previously performed a genome-wide screen to identify genes that function to maintain the stability of the rDNA region (13). In that screen, we used a collection of mutants in which each one of ∼4,800 non-essential genes of budding yeast is deleted, isolated the genomic DNA, and analyzed the size and size heterogeneity of chromosome XII of each mutant by pulsed-field gel electrophoresis (PFGE). The screen identified 708 mutants that showed instability of the rDNA region relative to wild-type (WT) cells (13). Many of the genes deleted in these mutants were found to function in nucleic acid transactions such as DNA replication, repair, and recombination. Unexpectedly, however, we also identified genes that had annotated functions in biological processes that seem to be unrelated to maintaining genome stability, such as cell and organelle morphogenesis (13). In this study, we sought to understand how these latter genes contribute to rDNA stability and found that Clb5 is a novel factor that is required for maintaining rDNA stability.

Eukaryotic genomes contain numerous replication origins that have the potential to fire but only a subset of them are activated to initiate DNA replication in a given S phase (14). The *CLB5* gene and its paralog *CLB6* encode S-phase cyclins, which regulate the timely activation of replication origins in non-rDNA regions in complex with cyclin-dependent kinase. Cells lacking Clb5 show prolonged S phase and defects in origin firing during late S phase, whereas absence of Clb6 has little effect on the duration of S phase or origin firing as long as Clb5 is present (15–18). In the *clb5 clb6* double mutant, entry into S phase is delayed but the length of S phase and origin firing are restored to WT levels (15–18). These observations suggest that Clb5 and Clb6 both promote timely activation of early firing origins, but only Clb5 is responsible for late origin firing.

Here we show that rDNA instability in the *clb5*Δ mutant is mostly suppressed when programmed replication fork arrest at the RFB site in rDNA is inhibited by *fob1* mutation. Deletion of the *CLB5* gene resulted in a reduction in the efficiency of replication origin firing in rDNA by half relative to WT cells. Both the rDNA instability and origin firing defects in *clb5*Δ cells were suppressed by the additional deletion of *CLB6*. The level of arrested forks was comparable in the tested strains, regardless of the presence or absence of Clb5. In the absence of Clb5, therefore, fewer replication forks initiate DNA replication in the rDNA and stall at the RFB, but these forks are more stably arrested because the arrival of the converging forks is delayed. Although the absence of Clb5 did not influence the level of DSBs or resected DSBs, it resulted in a higher level of recombination intermediates. Thus, persistently arrested forks at the RFB site may lead to DSB-independent, recombination-dependent rDNA instability. Alternatively, absence of Clb5 results in an increase in the distance between active origins due to inefficient replication initiation in rDNA, potentially leading to replication stress that causes damage at non-RFB sites and recombination-mediated rDNA instability.

## MATERIALS AND METHODS

### Yeast strains, growth conditions, and genomic DNA preparation

The mutant strains used in Figs. S1 and S2 are derivatives of the BY4741 background (*MATa his3*Δ*1 leu2*Δ*0 met15*Δ*0 ura3*Δ*0*) and obtained from the Yeast Knockout Collection (YSC1053; Open Biosystems [now at Horizon Discovery]) (19–21). The WT strain used in Figs. S1 and S2 was BY4741. The other strains used in this study were derived from NOY408-1b, which is in the W303 background (*MATa ade2-1 ura3-1 his3-11,15 trp1-1 leu2-3,112 can1-100*). Mutant strains in which genes of interest were deleted were constructed by a standard one-step gene replacement method, followed by PCR-based genotyping. The mutant strains used in Fig. 1 and Fig. S3 were constructed by replacing the ORF of the gene of interest with the kanMX marker in the NOY408-1b strain. The *clb5*Δ and *clb5*Δ *fob1* strains used in Fig. 2 were constructed by replacing the *CLB5* ORF with the kanMX marker in NOY408-1b and NOY408-1bf (NOY408-1b *fob1::LEU2*), respectively. The strains used in Figs. 3 and 4 and Fig. S4 were isolated by tetrad dissection of a diploid strain heterozygous for *fob1::LEU2, clb5*Δ::*kanMX*, and *clb6*Δ*::hphMX*, which was constructed by sequentially replacing the *CLB5* and *CLB6* ORFs with the kanMX and hphMX markers.

**Figure 2.**
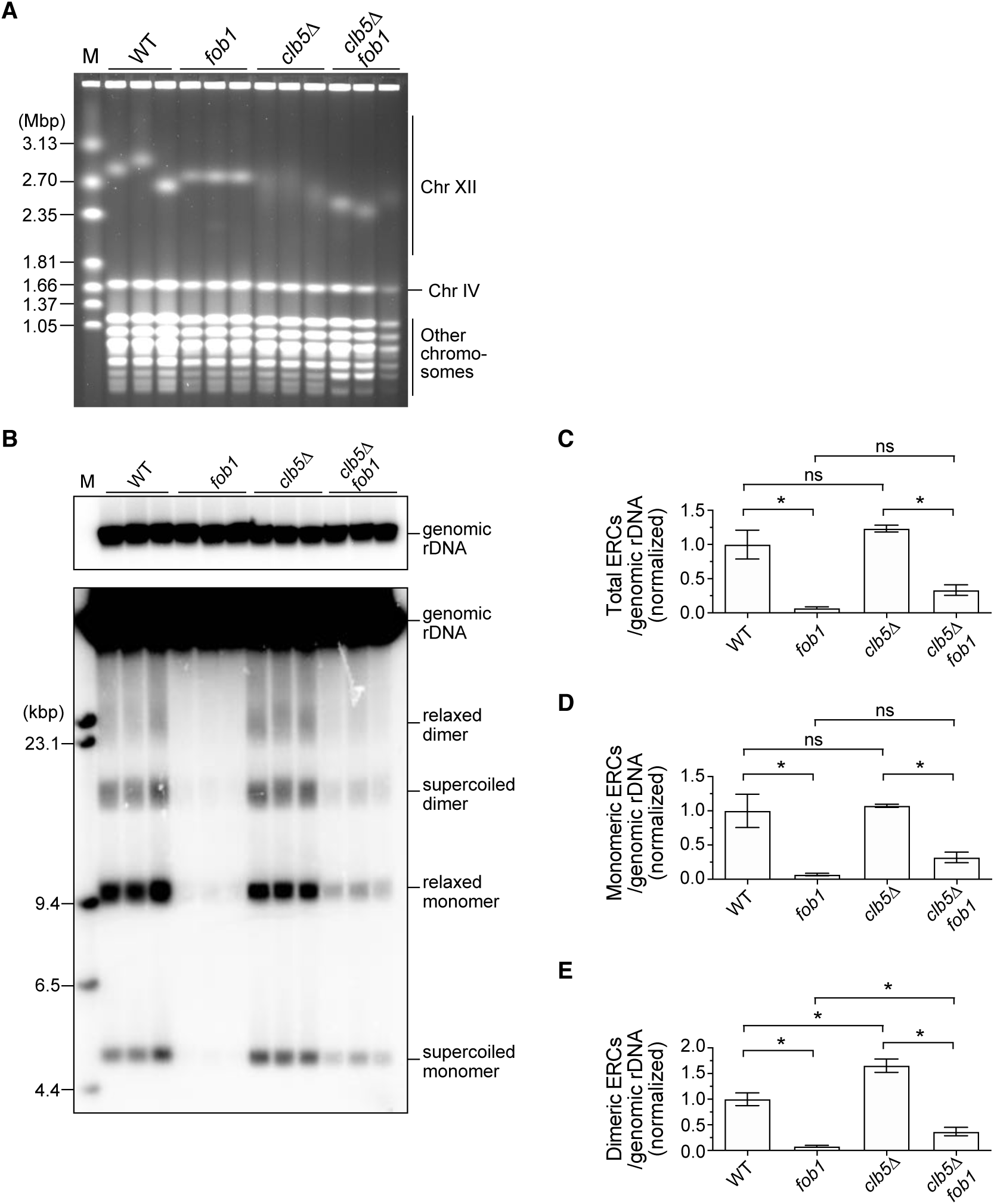
rDNA instability in the *clb5*Δ mutant is mainly dependent on Fob1. (A) PFGE analysis of the size heterogeneity of chromosome XII. DNA was extracted from three independent clones of the indicated strains and separated by PFGE. DNA was stained with ethidium bromide. M indicates *H. wingei* chromosomal DNA markers. (B) ERC detection. DNA was separated by conventional agarose gel electrophoresis, followed by Southern blotting with the rDNA probe. Genomic rDNA, and supercoiled and relaxed forms of monomeric and dimeric ERCs are indicated. M indicates λ DNA-*Hind* III markers. (C–E) Levels of total monomers and dimers (C), monomers (D), and dimers (E) relative to genomic rDNA. The level of ERCs in each mutant was normalized to the average of the WT clones (bars show mean ± SD). One-way ANOVA was used for multiple comparisons. Asterisks indicate a significant difference at p < 0.05; ns indicates that the difference is not significant (p >0.05).

**Figure 3.**
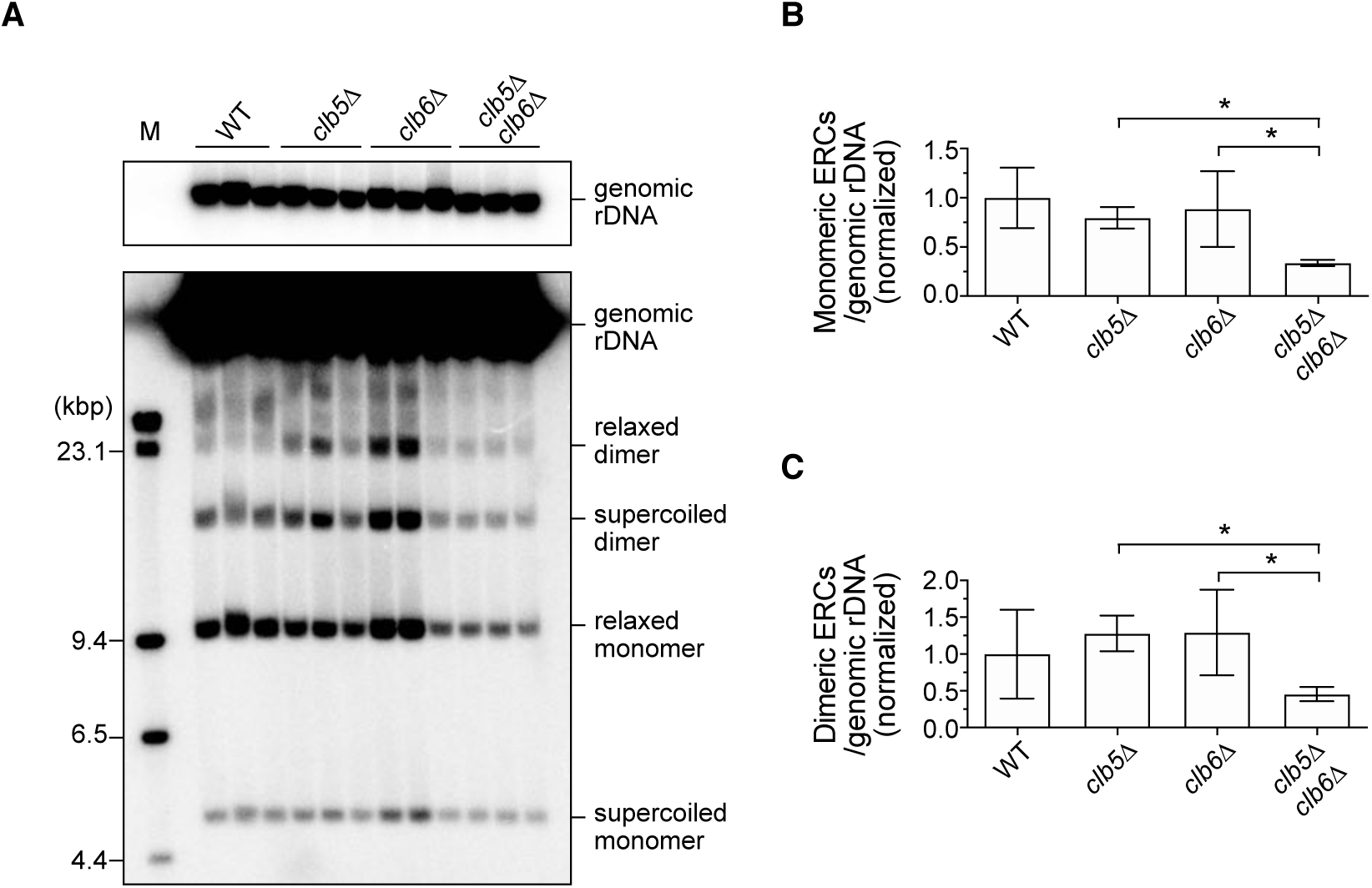
Production of ERCs in the *clb5*Δ mutant is suppressed by deletion of *CLB6*. (A) ERC detection. DNA was extracted from three independent clones of the indicated strains and separated by conventional agarose gel electrophoresis, followed by Southern blotting with the rDNA probe. Genomic rDNA, and supercoiled and relaxed forms of monomeric and dimeric ERCs are indicated. M indicates λ DNA-*Hind* III markers. (B, C) Levels of monomers (B) and dimers (C) relative to genomic rDNA. The level of ERCs in each mutant was normalized to the average of WT clones (bars show mean ± SD for six independent clones). One-way ANOVA was used for multiple comparisons. Asterisks indicate a significant difference at p < 0.05.

**Figure 4.**
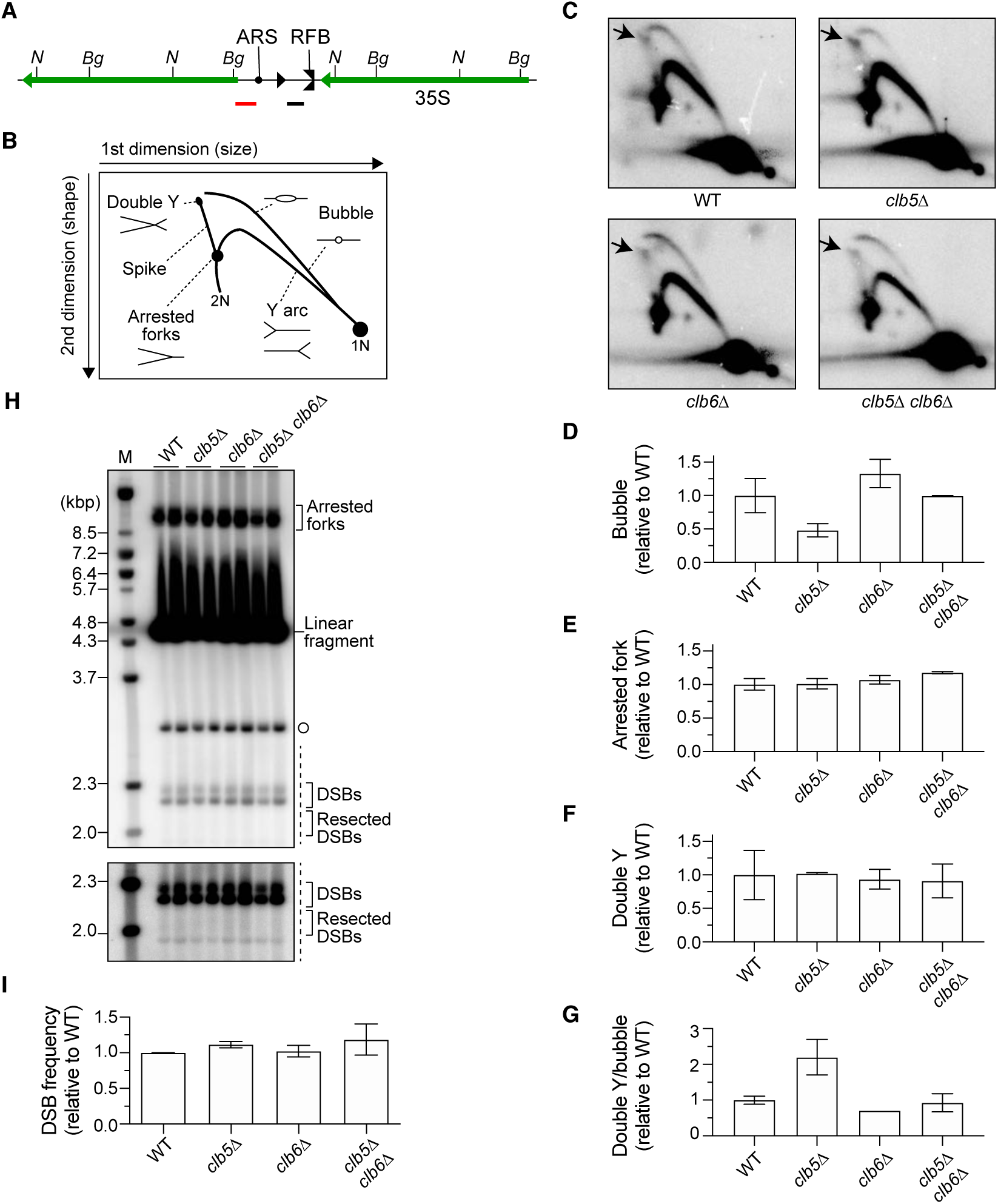
Frequencies of replication initiation, replication fork arrest, and DSBs in *clb5*Δ and *clb6*Δ mutants. (A) Restriction map of the rDNA repeat. Organization of the rDNA repeats is indicated as in Fig. 1A. *N* and *Bg* indicate the sequences recognized by restriction enzymes *Nhe* I and *Bgl* II, respectively. Black and red bars represent the Southern blotting probes used for 2D and DSB analyses, respectively. (B, C) 2D agarose gel electrophoresis. Genomic DNA was digested with *Nhe* I. DNA was separated by size in the first dimension and by size and shape in the second dimension, and subjected to Southern blotting with the rDNA probe indicated in (A). The expected migration pattern of different replication intermediates by 2D analysis is shown in (B). 1N and 2N indicate, respectively, one- and two-unit length linear DNA molecules. Arrows in (C) indicate a double Y spot. (D, E, F) Frequency of bubble arcs (D), arrested forks (E), and double Y spots (F) determined by quantifying the signal of each intermediate relative to the total amount of replication intermediates. The levels of the different molecules in each mutant were normalized to the average of WT clones (bars indicate the range of two independent experiments). (G) Frequency of recombination intermediates at the RFB site determined by the ratio of the double Y spot signal to the bubble arc signal, which was normalized to the average of WT clones (bars indicate the range of two independent experiments). (H) Detection of arrested forks and DSBs. Genomic DNA was digested with *Bgl* II and separated by single-dimension agarose gel electrophoresis, followed by Southern blotting with the rDNA probe indicated in (A). Bands corresponding to arrested forks, linear fragments, DSBs, and resected DSBs are indicated. Open circle represents the terminal fragment containing the telomere-proximal rDNA repeat and its adjacent non-rDNA fragment. The lower panel shows a more exposed contrast image of the phosphorimager signal in the region marked by the dashed line. M indicates λ DNA-*BstE* II markers. (I) Frequency of DSBs determined as the ratio of DSB signal to arrested fork signal, which was normalized to the average of WT clones (bars indicate ranges of two independent experiments).

For the PFGE and extrachromosomal rDNA circle (ERC) analyses in Figs. S1 and S2, yeast strains were patched from their glycerol stock onto YPD plates (1% [w/v] yeast extract, 2% [w/v] peptone, 2% [w/v] glucose, and 2% [w/v] agar) and the bulk of cells were grown in 5 ml of YPD medium overnight at 30°C. For other PFGE and ERC analyses, yeast strains were streaked onto YPD plates, a single colony was then inoculated into 5 ml of YPD medium, and cells were grown overnight at 30°C. Cells (5 x 10^7^ cells/plug) were collected and washed twice with 50 mM EDTA (pH 7.5).

For cells subjected to two-dimensional (2D) and DSB analyses, a single colony was inoculated into 5 ml of YPD medium and grown overnight at 30°C until the culture reached the saturation phase of growth. The cells were then inoculated into 100 ml of YPD medium at an OD_600_ of 0.1 and grown at 30°C until an OD_600_ of 0.4. The cells were immediately treated with 0.1% sodium azide and then collected (5 x 10^7^ cells/plug) and washed twice with 50 mM EDTA (pH 7.5). For PFGE, ERC, 2D, and DSB analyses, genomic DNA was prepared in low-melting-temperature agarose plugs as described previously (22).

### PFGE analysis

One-third of an agarose plug, along with *Hansenula wingei* chromosomal DNA markers (Bio-Rad), was separated by electrophoresis on a 1.0% agarose gel (pulsed-field certified agarose, Bio-Rad) in 0.5× Tris-borate EDTA (TBE) buffer (44.5 mM Tris base, 44.5 mM boric acid, and 1 mM EDTA, pH 8.0) on a Bio-Rad contour-clamped homogeneous electric field DR-III system using the following conditions: 68 hr at 3.0 V/cm, 120° included angle, linear ramp from 300 s of initial switch time to 900 s of final switch time. The gel was stained with 0.5 μg/ml of ethidium bromide and photographed.

### ERC assay

One-half of an agarose plug, along with 500 ng of Lambda *Hind* III DNA markers, was separated by electrophoresis on a 0.4% agarose gel (15 × 25 cm gel) in 1× Tris-acetate-EDTA (40 mM Tris base, 20 mM acetic acid, and 1 mM EDTA, pH 8.0) at 1.0 V/cm for ∼48 hr at 4°C with buffer circulation in a Sub-cell GT electrophoresis system (Bio-Rad). The buffer was changed every ∼24 hr.

DNA was transferred to Hybond XL (GE Healthcare). Southern blotting was then performed with a probe prepared by PCR amplification of genomic DNA using primers 5′-CATTTCCTATAGTTAACAGGACATGCC and 5′-AATTCGCACTATCCAGCTGCACTC, as described previously (22). The membrane was exposed to phosphor screens for an appropriate amount of time before any signals were saturated, and the radioactive signal was detected using Typhoon FLA7000 (GE Healthcare). The membrane was re-exposed to the phosphor screen for several days and scanned. The scanned images taken after short and long exposures were used to quantify genomic rDNA and ERC bands, respectively, using FUJIFILM Multi Gauge version 2.0 software (Fujifilm). The ratio of ERCs relative to genomic rDNA was determined.

### 2D gel electrophoresis

2D gel electrophoresis was performed as described previously with slight modifications (23). In brief, one-half of an agarose plug was placed in a 2-ml flat-bottom tube. The plug was equilibrated twice in 1 ml of 1× M buffer (TaKaRa) by rotating the tube for 30 min at room temperature. After discarding the buffer completely, the plug was incubated in 160 μl of 1× M buffer containing 160 units of *Nhe* I (TaKaRa) for 7 hr at 37°C. The plug and 600 ng of Lambda *Hind* III DNA markers were separated by electrophoresis on a 0.4% agarose gel (SeaKem Agarose LE, Lonza) in 1× TBE buffer at 1.32 V/cm for 14 hr at room temperature with buffer circulation in a Sub-cell GT electrophoresis system (15 × 20 cm gel, Bio-Rad). The gel was stained with 1× TBE buffer containing 0.3 μg/ml of ethidium bromide and photographed. Gel slices containing DNA of the size ranging from 4.7 to 9.4 kb were excised, rotated 90°, and cast in a 1.2% agarose gel (SeaKem Agarose LE, Lonza) containing 0.3 μg/ml of ethidium bromide in 1× TBE. The second-dimension gel electrophoresis was performed in 1× TBE buffer containing 0.3 μg/ml of ethidium bromide at 6.0 V/cm for 5 hr at 4°C with buffer circulation in a Sub-cell GT electrophoresis system (Bio-Rad). DNA was transferred to Hybond-XL (GE Healthcare).

Southern blotting was performed with a probe prepared by PCR amplification of genomic DNA using primers 5′-CATTTCCTATAGTTAACAGGACATGCC and 5′-AATTCGCACTATCCAGCTGCACTC, as described previously (22). The membrane was exposed to phosphor screens for several days and the radioactive signal was detected using Typhoon FLA7000 (GE Healthcare). ImageJ (NIH) was used to quantify bubbles, Y arcs containing RFB spots, RFB spots, and double Y spots.

### DSB assay

The DSB assay was performed as described previously (22). In brief, one-third of an agarose plug was placed in a 2-ml flat-bottom tube. The plug was equilibrated four times in 1 ml of 1× TE (10 mM Tris base, pH 7.5, and 1 mM EDTA, pH 8.0) by rotating the tube for 15 min at room temperature. The plug was then equilibrated twice in 1 ml of 1× NEBuffer 3.1 (New England Biolabs) by rotating the tube for 30 min at room temperature. After discarding the buffer completely, the plug was incubated in 160 μl of 1× NEBuffer 3.1 buffer containing 160 units of *Bgl* II (New England BIolabs) overnight at 37°C. The plug and 600 ng of Lambda *Bst* EII DNA markers were separated by electrophoresis on a 0.7% agarose gel (15 × 20 cm gel) in 1× TBE buffer at 2.0 V/cm for 21 hr at room temperature with buffer circulation in a Sub-cell GT electrophoresis system (Bio-Rad). The gel was stained with 0.5 μg/ml of ethidium bromide.

DNA was transferred to Hybond XL. Southern blotting was performed with a probe prepared by PCR amplification of genomic DNA using primers 5′-ACGAACGACAAGCCTACTCG and 5′-AAAAGGTGCGGAAATGGCTG, as described previously (22). The membrane was exposed to phosphor screens overnight. The radioactive signal was detected using Typhoon FLA7000 (GE Healthcare). Signals of DSBs and arrested forks were quantified using FUJIFILM Multi Gauge version 2.0 software (Fujifilm). The ratio of DSBs relative to arrested forks was determined and normalized to the average value of WT samples.

### Statistical analysis

Statistical analysis was performed by using GraphPad Prism software (version 8.0).

## RESULTS

### The rDNA region is destabilized in the absence of Clb5

We previously identified 708 candidate genes that may contribute to maintaining rDNA stability and conducted Gene Ontology analysis (13). We have tested the reproducibility of rDNA instability in some mutants lacking genes known to function in nucleic acid metabolism such as DNA replication, recombination, repair, and transcription (13, 22, 24–26). Among the original 708 rDNA-unstable mutants, however, 113 mutants lack a gene that has an annotated function seemingly unrelated to genome stability but instead is involved in biological processes such as membrane invagination, endocytosis, cytokinesis, nucleus organization, conjugation, cell morphogenesis, cell wall organization or biogenesis, cytoskeleton organization, sporulation, membrane fusion, pseudohyphal growth, invasive growth in response to glucose limitation, and exocytosis or cell budding (13).

In our previous genome-wide screen, we assessed the degree of rDNA stability of each of ∼4,800 mutants only once by PFGE, because the number of mutants was too large to analyze in multiplicate. Furthermore, we used a comb with the thinnest teeth to improve throughput, which might have reduced the sharpness of the band and produced false-positives, as discussed previously (27). In addition, we now possess an updated version of the original yeast deletion collection. Here, therefore, we first confirmed the reproducibility of rDNA instability in candidate mutants with a GO annotation unrelated to genome stability that were directly obtained from the updated yeast deletion collection. Because 15 of the 113 mutants were analyzed in our previous study (26), and 1 mutant could not be revived from the glycerol stock, the remaining 97 mutants were subjected to confirmation for their rDNA instability.

To assess the rDNA instability, we isolated genomic DNA from the candidate mutants and separated DNA by PFGE using a comb with wider teeth as compared with our previous screen to improve the sensitivity of our analysis. We then examined the size heterogeneity of chromosome XII, which carries the rDNA array (Fig. S1). The *SIR2* gene encoding a histone deacetylase is known to promote rDNA stability (10, 28, 29); thus, in each gel, we included DNA from WT and the *sir2*Δ mutant, which exhibits an extremely smeared band of chromosome XII as compared with WT cells (Fig. S1). PFGE analysis showed that the chromosome XII band was prominently smeared in 15 mutants: *ypt7*Δ (Fig. S1A), *bud6*Δ, *svl3*Δ, *clb5*Δ, *rho4*Δ, *pam1*Δ, *swm1*Δ, *pep7*Δ (Fig. S1B), *hcm1*Δ, *ldb17*Δ, *siw14*Δ (Fig. S1C), *bem2*Δ, *vrp1*Δ, *pho85*Δ, and *gon7*Δ (Fig. S1D).

Changes in rDNA copy number are often associated with the production of extrachromosomal rDNA circles (ERCs), which are excised from genomic arrays. Many of the previously studied rDNA-unstable mutants, such as the *sir2*Δ mutant, accumulate ERCs at a level higher than that of WT cells (28, 29). To examine the level of ERCs in the candidate mutant strains, we separated genomic DNA by conventional agarose gel electrophoresis to resolve ERCs from genomic rDNA, and then detected ERCs and genomic rDNA by Southern blotting (Fig. S2). As a reference, genomic DNA samples from WT and the *sir2*Δ mutant were included in the analyses. ERCs were not clearly detected in one of the genomic DNA samples prepared from WT cells due to poor sample quality (Fig. S2), and faint ERC bands were detected in the other WT sample. Therefore, we did not use the WT sample as the control, but instead compared the ERC level in each mutant to that in the *sir2*Δ mutant as a positive control. The above-mentioned mutants that exhibited a prominently smeared band of chromosome XII in PFGE did not accumulate ERCs to a level comparable to that in the *sir2*Δ mutant (Figs. S1 and S2). Instead, *vac8*Δ, *chs7*Δ, *cdc10*Δ, *hub1*Δ, and *rim11*Δ were the highest-ranked mutants and produced ERCs at a level comparable to that in *sir2*Δ, although they did not show severe rDNA instability in PFGE analysis (Figs. S1 and S2). Collectively, these analyses enabled us to narrow down the number of genes to test further for confirmation of rDNA instability.

To examine the involvement of these 20 candidate genes in maintaining rDNA stability, we constructed deletion mutants of each gene in the W303 yeast strain background that differs from the BY background in which the yeast deletion collection was generated. For unknown reasons, it was not possible to construct mutants lacking *CDC10, PAM1, PEP7, PHO85*, or *GON7*. Among the remaining 15 mutants, only *clb5*Δ showed a smeared band of chromosome XII (Fig. 1B and 1C). In this mutant, however, the ERC level was similar to that in WT cells (Fig. S3C and S3D). None of the other 14 mutants produced ERCs at a level comparable to that in the *sir2*Δ mutant (Fig. S3). Taken together, these findings show that absence of Clb5 causes severe rDNA instability in yeast strains of two different genetic backgrounds, demonstrating that Clb5 plays an important role in maintaining rDNA stability.

A previous study showed that the *clb5*Δ mutant exhibits small defects in vacuolar fragmentation under various growth conditions (30). GO analysis thus associates the *CLB5* gene with the biological processes of organelle assembly, organelle fission, and regulation of organelle organization, in addition to its best-known function as an S-phase cyclin required for the activity of CDK, which regulates the initiation and progression of DNA replication (15–18).

### Clb5 suppresses rDNA instability in response to Fob1-mediated replication fork arrest

To understand how Clb5 promotes rDNA stability, we examined whether rDNA instability in the *clb5*Δ mutant occurs in response to Fob1-mediated replication fork arrest at the RFB. To this end, we compared rDNA stability in WT, *fob1, clb5*Δ, and *clb5*Δ *fob1* cells by PFGE. Smearing of the chromosome XII band in the *clb5*Δ mutant was largely suppressed by the introduction of a *fob1* mutation (Fig. 2A). The chromosome XII band in the *clb5*Δ *fob1* mutant, however, was slightly broader than that in the *fob1* single mutant (Fig. 2A). These results suggest that rDNA instability in the *clb5*Δ mutant mostly depends on Fob1.

We also assessed rDNA instability by ERC analysis (Fig. 2B–2E). The electrophoresis conditions used allowed the detection of super-coiled and relaxed forms of monomeric and dimeric ERCs (Fig. 2B). The total level of monomers and dimers in the *clb5*Δ mutant was comparable to that in WT (Fig. 2C), as observed in Fig. S3C and S3D. When we analyzed monomeric and dimeric ERCs separately, the level of dimeric ERCs was increased in *clb5*Δ by 1.7-fold relative to WT (Fig. 2E), whereas that of monomeric ERCs was similar between WT and *clb5*Δ (Fig. 2D). Thus, rDNA instability in the absence of Clb5 seems to be associated with the production of multimeric ERCs.

Consistent with our PFGE analysis demonstrating that rDNA instability in the *clb5*Δ mutant was mostly suppressed by *fob1* mutation (Fig. 2A), the level of monomeric and dimeric ERCs was substantially reduced by 3.3- and 4.5-fold, respectively, in the *clb5*Δ *fob1* mutant relative to the *clb5*Δ mutant (Fig. 2D, 2E). The level of dimeric ERCs in the *clb5*Δ *fob1* mutant, however, was still higher than that in the *fob1* mutant (Fig. 2E). The *clb5*Δ *fob1* mutant also accumulated more monomeric ERCs as compared with the *fob1* mutant, but the difference was not statistically significant (Fig. 2D). Thus, in the absence of Clb5, ERCs are produced predominantly in a Fob1-dependent manner, but some are generated independently of Fob1. Taken together, these results suggest that Clb5 functions to maintain rDNA instability mainly by promoting the proper response to Fob1-mediated replication fork arrest, but it may also prevent the generation of rDNA-destabilizing DNA damage in non-RFB regions.

### rDNA instability in the *clb5*Δ mutant is suppressed by deletion of its paralog *CLB6*

*CLB5* has a paralog, *CLB6*, whose product also acts as an S-phase cyclin. Whereas deletion of *CLB5* causes lengthening of S phase, deletion of *CLB6* has little effect on S-phase duration (16–18). In the *clb5 clb6* double mutant, the onset of S phase is delayed, but the length of S phase is restored to that seen in WT (18). Thus, Clb5 and Clb6 differentially regulate the initiation and progression of S phase.

Next, we sought to understand how Clb5 and Clb6 coordinately regulate rDNA instability. There were no differences in the level of monomeric ERCs among WT, *clb5*Δ and *clb6*Δ cells (Fig. 3A, 3B). The *clb5*Δ mutant showed a higher level of dimeric ERCs as compared with WT, although the difference was not statistically significant this time (Fig. 3C). The production of monomers and dimers in the *clb5*Δ and *clb6*Δ mutants was suppressed by 2.4- to 2.8-fold in the *clb5*Δ *clb6*Δ mutant (Fig. 3B, 3C). PFGE analysis also showed similar patterns in which smearing of the chromosome XII band appeared less severe in the *clb5*Δ *clb6*Δ double mutant than in the *clb5*Δ single mutant (Fig. S4). Thus, rDNA instability in the *clb5*Δ mutants is suppressed by deletion of *CLB6*.

### Clb5 regulates replication initiation in the rDNA region

Previous studies have demonstrated that Clb5 and Clb6 differentially regulate the timing of replication origin firing in non-rDNA regions: that is, Clb5 and Clb6 both promote firing of early replication origins, but only Clb5 activates late origins (15, 18). Absence of both Clb5 and Clb6 delays entry into S phase, but restores the length of S phase and origin firing to WT levels (15–18). To understand how Clb5 and Clb6 regulate replication in the rDNA region, we assessed the frequency of origin firing in this region in the *clb5*Δ and *clb6*Δ mutants (Fig. 4A–4D).

To this end, we first grew cells to stationary phase, diluted them into fresh media, and allowed them to grow until they re-entered the cell cycle, enabling us to obtain a sample enriched in replicating cells. We then prepared genomic DNA, digested it with the restriction enzyme *Nhe* I, which has two recognition sites within each rDNA unit (Fig. 4A), and performed two-dimensional (2D) agarose gel electrophoresis. DNA molecules were separated by molecular mass in the first dimension, followed by mass and shape in the second dimension. DNA was transferred to a nylon membrane and analyzed by Southern blotting with a probe that hybridizes to the *Nhe* I fragment containing an origin of DNA replication and an RFB site (Fig. 4A).

In this analysis, DNA molecules where replication is initiated from an origin are detected as bubbled-shaped molecules, while those where replication passively progresses through the *Nhe* I fragment generate a Y arc (Fig. 4B) (31). In addition, replication forks that are paused at the RFB site within the restriction fragment produce an intensive signal along the Y arc (Fig. 4B). We assessed the frequency of origin firing by determining the proportion of signal for bubbled-shaped molecules relative to the total signal for replication intermediates (Fig. 4B, 4C). The efficiency of origin firing in the *clb5*Δ mutant was lowered by half relative to WT (Fig. 4D), demonstrating that Clb5 promotes firing of replication origins in the rDNA. The reduced origin firing observed in the *clb5*Δ mutant was restored in the *clb5*Δ *clb6*Δ double mutant to almost WT levels (Fig. 4D). Together with the finding that rDNA instability in the *clb5*Δ mutant was suppressed in the *clb5*Δ *clb6*Δ mutant (Fig. 3), these results indicate that rDNA stability is influenced by the efficiency of replication initiation.

### Replication forks are more stably arrested at the RFB site in the absence of Clb5

To understand how reduced origin firing in the *clb5*Δ mutant leads to rDNA instability, we examined whether absence of Clb5 influences the frequency of replication fork arrest. More than 90% of replication forks that initiate from replication origins stall at the RFB site (9). Because fewer origins fired in the absence of Clb5 (Fig. 4D), we expected that fewer forks would be stalled at the RFB site in the *clb5*Δ mutant as compared with WT cells. We assessed the frequency of arrested forks by determining the proportion of signal for arrested fork intermediates relative to the total signal for replication intermediates in our 2D analysis (Fig. 4B, 4C). The fork blocking activity was comparable among WT, *clb5*Δ, *clb6*Δ, and *clb5*Δ *clb6*Δ cells (Fig. 4E). This finding implies that, although fewer forks are stalled at the RFB in the *clb5*Δ mutant compared with WT cells, these forks are more stably arrested, resulting in a higher level of persistently arrested forks.

Replication fork arrest at the RFB site leads to the formation of DSBs, which are thought to be a major trigger of genome instability (2, 32). It had long been thought that DSBs formed at arrested forks are repaired by homologous recombination; however, our previous study demonstrated that DSBs are rarely resected and their repair does not require recombination proteins in WT cells as long as the cells carry a normal rDNA copy number (22). Thus, changes in rDNA copy number can be dictated by the frequency of DSB end resection.

To determine the frequency of DSBs and resected DSBs formed at the RFB site, we digested DNA with the restriction enzyme *Bgl* II, and separated it by single-dimension agarose gel electrophoresis, followed by Southern blotting (Fig. 4A, 4H). We assessed the DSB frequency by determining the ratio of signal for DSBs to that for arrested fork intermediates (Fig. 4H, 4I). DSB levels were comparable among WT, *clb5*Δ, *clb6*Δ and *clb5*Δ *clb6*Δ cells (Fig. 4I). Resected DSBs in the *clb5*Δ mutant were below the detection limit of Southern blotting, but this result indicates that absence of Clb5 does not seem to cause a substantial increase in DSB end resection (Fig. 4H). Thus, Clb5 does not influence the formation or repair of DSBs.

### The *clb5*Δ mutant accumulates recombination intermediates

In 2D gel analysis, we also detected replication forks converging from both sides at the RFB as well as Holliday junction recombination intermediates that are formed during homologous recombination-mediated repair of DSBs (31). Both intermediates generate a double Y spot signal on the top of spike signal and were seen in a similar position in our 2D gel (Fig. 4B, 4C). As shown in Fig. 4C, the double Y spot signal in the *clb5*Δ mutant seemed to be stronger than that in WT, *clb6*Δ, or *clb5*Δ *clb6*Δ cells. However, the double Y spot signal, as a proportion of total replication intermediate signals, was similar among these cells (Fig. 4F), possibly because more replication intermediates may exist in *clb5*Δ, which has a longer S phase. In contrast, the ratio of double Y spot to bubble signals was clearly increased in the *clb5*Δ mutant (Fig. 4G). Because the arrested fork signal was not elevated (Fig. 4E), and the bubble signal was reduced (Fig. 4D) in *clb5*Δ relative to WT, *clb6*Δ, and *clb5*Δ *clb6*Δ cells, the increased signal ratio of double Y spots in the *clb5*Δ mutant is most probably due to a higher level of recombination intermediates but not converging replication forks. Collectively, these results suggest that recombination frequency is increased in the *clb5*Δ mutant, possibly leading to rDNA instability.

## DISCUSSION

In this study, we confirmed the reproducibility of rDNA instability in 97 previously identified rDNA-unstable mutants that lack genes with a GO annotation that is seemingly unrelated to genome stability—for example, genes functioning in cell and organelle morphogenesis (13). We demonstrated that a mutant lacking the *CLB5* gene exhibits rDNA instability. Based on GO analysis, *CLB5* is associated with the biological processes of organelle fission and regulation of organelle organization, because its deletion causes small defects in vacuolar fragmentation under various growth conditions (30). However, the mechanism by which *CLB5* regulates vacuolar fragmentation remains unknown. The *CLB5* gene encodes an S-phase cyclin required for the activity of Cdc28 and is involved in the initiation and progression of DNA replication (15–18). In this study, we have revealed that Clb5 plays an important role in suppressing rDNA instability, mainly in response to Fob1-mediated replication fork arrest and partially in response to DNA damage at non-RFB sites, by promoting replication initiation in the rDNA region.

The initiation of replication across the genome is under temporal control. In non-rDNA regions, Clb5 and Clb6 function redundantly to promote the timely activation of early firing origins, but only Clb5 activates late origins (15). In the rDNA region, each rDNA copy contains a replication origin sequence with the potential to initiate replication, but only a small proportion of these origins fire during rDNA replication (6). We found that deletion of the *CLB5* gene results in a 50% reduction in replication initiation efficiency during exponential growth as compared with WT cells, whereas deletion of *CLB6* has little effect on replication initiation (Fig. 4D). Thus, Clb5 plays a dominant role in the activation of replication origins in the rDNA region. Although we did not examine the timing of origin activation, we speculate that, in the absence of Clb5, late origin firing is defective in the rDNA region (Fig. 5).

**Figure 5.**
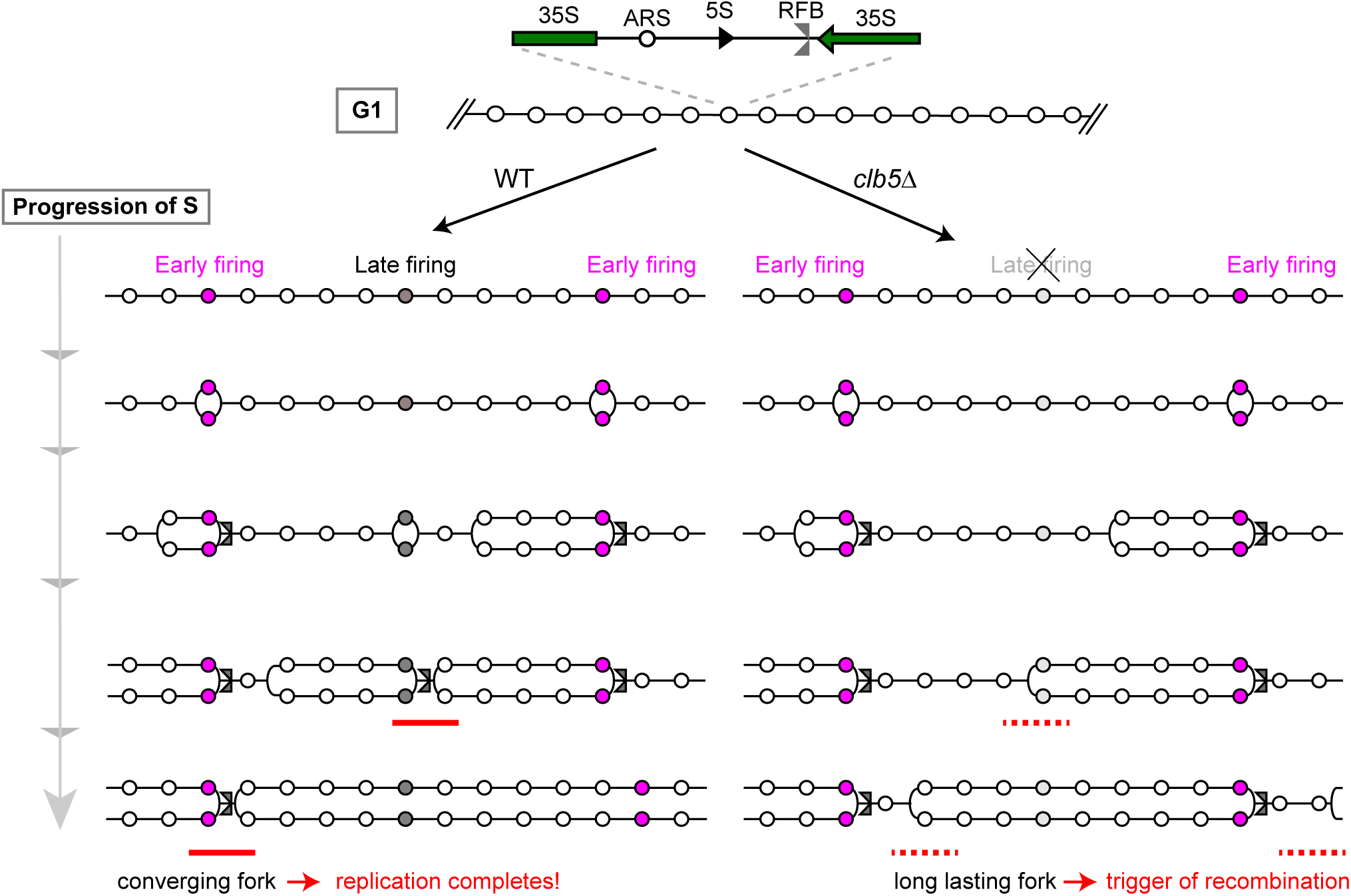
Model of rDNA replication in a replication initiation-reduced condition. (Top) Part of rDNA in G1 phase. Circles represent replication origins (ARS). (Left) Progression of DNA replication in WT. The pink-shaded ARSs start replication in early S phase, while the gray-shaded ARSs start late. Right-ward moving forks are arrested by the function of Fob1 at the RFB site. About one in five ARSs starts replication in S-phase (6, 35). The red line indicates a converging fork. (Right) Progression of DNA replication in the *clb5*Δ mutant. Right-ward moving forks are arrested by the function of Fob1 at the RFB site and remain arrested until the converging fork arrives. Left-ward moving forks have to travel a longer distance because there are fewer late-firing origins. The dashed red line indicates a long-lasting fork. The number of RFB-arrested forks is similar in WT and *clb5*Δ (six are shown), but there are fewer converging forks in the *clb5*Δ mutant. As a result, replication forks in the *clb5*Δ mutant have a longer distance to travel and may encounter problems, such as accidental fork arrest that triggers recombination, on the way.

We previously demonstrated that the proper regulation of replication initiation is important to maintain rDNA stability (33). When the rDNA copy number is reduced to less than half (∼80) of the normal number, cells undergo rDNA expansion until they reach the normal copy number in a process referred to as “gene amplification” (34). The rate of rDNA amplification is reduced in cells carrying a modified form of the replication origin sequence rARSΔ-3, which exhibits lower replication initiation activity in the rDNA region (33). Our observation of rDNA instability in the *clb5*Δ mutant is consistent with that previous finding, although the rARSΔ-3 strain does not accumulate ERCs (discussed below). Previous studies demonstrated that the prolonged S phase and late origin firing defects in the non-rDNA regions seen in the *clb5*Δ mutant are suppressed in the *clb5*Δ *clb6*Δ mutant, although the double mutant has delayed entry into S phase (15, 18). It has been proposed that mitotic cyclins may promote S phase in the absence of both Clb5 and Clb6 (15). The rDNA instability and replication initiation defect observed in the *clb5*Δ mutant were also suppressed in the *clb5*Δ *clb6*Δ mutant (Figs. 3 and 4), thereby suggesting that Clb6 either represses late origin firing or is not sufficient to activate late origins in the rDNA region in the absence of Clb5. These findings demonstrate that the proper activation of late firing origins may be critical for maintaining rDNA stability.

Relative to WT cells, the *clb5*Δ mutant showed a higher level of multimeric ERCs (Fig. 2E), whereas the rARSΔ-3 strain shows a lower level of ERCs (33). The reason behind these differences in ERC levels remains unclear. We envisage that, in the rARSΔ-3 strain, the efficiency of replication initiation is compromised for both early and late firing origins, while the timing of their activation remains unaltered. In the *clb5*Δ mutant, by contrast, early firing origins are activated at the normal time but late firing origins are inefficiently activated, raising the possibility that ERCs may be produced by recombination-related events that are associated with replication forks initiated in late S phase.

Once replication is initiated, the majority of replication forks (>90%) are immediately stalled at the nearest RFB site in a polar fashion (7, 9, 35). Thus, we expected that the *clb5*Δ mutant, which has fewer active origins, would generate fewer forks arrested at the RFB site. Unexpectedly, however, the level of arrested forks in the mutant lacking *CLB5* was comparable to that in WT cells (Fig. 4E), indicating that replication forks are more stably stalled at the RFB in the absence of Clb5. Previous studies have shown that active replication origins in the rDNA are clustered and separated by large gaps without active origins (36–38). A stalled replication fork can be resolved by the arrival of the converging fork proceeding from the nearest origin (3). For the *clb5*Δ mutant, which has fewer firing origins, the converging fork must come from an origin that is further away (Fig. 5). In the absence of Clb5, therefore, more forks remain arrested for a longer time, which is the initiating event for RFB-dependent rDNA instability (Fig. 5).

The replication termination sequence RTS1 induces site-specific fork stalling in fission yeast (1). Previous studies have demonstrated that genome rearrangements are induced by homologous recombination-mediated replication fork restart at stalled forks at the RTS1 site when the arrival of the converging fork is inhibited by the presence of another RTS1 site, but this process does not involve DSB formation (39, 40). It remains unknown whether replication forks arrested at the RFB site in budding yeast rDNA undergo similar genome rearrangements induced by replication fork restart. Nonetheless, absence of Clb5 delays the arrival of the converging fork by impairing the firing of upstream late origins and, as a result, the forks arrested at the RFB site persist longer (Fig. 5). These forks may undergo DSB-independent faulty template switches to restart DNA replication, leading to rDNA instability.

The other consequence of persistent replication fork arrest is the generation of DSBs, which have the potential to induce genome rearrangements when not repaired properly by equal sister-chromatid exchange (41). We found, however, that the level of DSBs at the RFB site was comparable, regardless of the presence or absence of Clb5 (Fig. 4I). Several factors can influence the outcome of DSB repair. First, one of the intergenic regions of the rDNA unit contains an RNA polymerase II-dependent, bi-directional promoter, named E-pro, upstream of the RFB site (42), and transcription of non-coding RNA from E-pro may induce the dissociation of cohesin from rDNA repeats, leading to unequal sister-chromatid recombination that alters the rDNA copy number during DSB repair (10, 42). Thus, Clb5 may suppress rDNA instability by regulating the transcription of non-coding RNA from E-pro and/or cohesin association—two possibilities that need to be examined in future studies. The occurrence of rDNA instability may also depend on the decision of whether cells undergo DSB end resection and repair by homologous recombination (22). In both WT and *clb5*Δ, resected DSBs were below the detection limit of our Southern blotting analysis (Fig. 4H). Therefore, it seems unlikely that absence of Clb5 induces DSB end resection at the RFB.

In our 2D analysis, the double Y spot signal was stronger in the *clb5*Δ mutant than in WT, *clb6*Δ, or *clb5*Δ *clb6*Δ cells (Fig. 4C and 4G). As mentioned above, this signal most probably corresponds to recombination intermediates. Therefore, DNA damage might occur more frequently in *clb5*Δ than in the other strains, which might explain why only *clb5*Δ shows rDNA instability among the mutants tested. If this is the case, what increases the damage in the *clb5*Δ mutant? Because the signals for arrested forks, DSBs, and resected DSBs at the RFB site were not increased in the *clb5*Δ mutant (Fig. 4E, 4H, 4I), we speculate that DNA damage occurs at non-RFB sites in the rDNA region in *clb5*Δ. In fact, although in PFGE the chromosome XII band in the *clb5*Δ *fob1* double mutant was sharper than that in the *clb5*Δ single mutant, it was still a little broader than that in the *fob1* single mutant (Fig. 2A). Moreover, in the ERC assay, more ERCs were detected in the *clb5*Δ *fob1* double mutant than in the *fob1* mutant (Fig. 2B–2E). These results suggest that some *FOB1* (RFB)-independent recombination occurs in *clb5*Δ. One possibility for triggering this recombination might be DNA damage caused by the reduced initiation of replication in the *clb5*Δ mutant (Fig. 5). It is known that a longer distance between replication origins induces more genome instability, making a site fragile (43). It has been speculated that the long-lasting forks have more time to cause problems, such as accidental fork arrest, during the course of travel between origins. A similar situation may occur in the rDNA of the *clb5*Δ mutant. Moreover, a long-lasting fork is expected to make bigger replication bubbles. This might enhance unequal-sister chromatid recombination, which would contribute to the rDNA copy number alteration and multimer ERC production seen in the *clb5*Δ mutant (Fig. 2). Further analysis is required to reveal the details of how long-lasting forks cause problems and their resulting DNA damage.

## ACKNOWLEDGMENTS

We thank members of the Laboratory of Genome Regeneration, especially Dr. Tetsushi Iida for discussion. This work was supported by JST CREST (Grant Number JP19207241 to T.K.) and in part by grants-in-aid for Scientific Research (17H01443 to T.K., and 18H04709 and 20H05382 to M.S.) from the Japan Society for the Promotion of Science. We have no conflicts of interest to declare.

## FIGURE LEGENDS

**Supplemental Figure 1.**
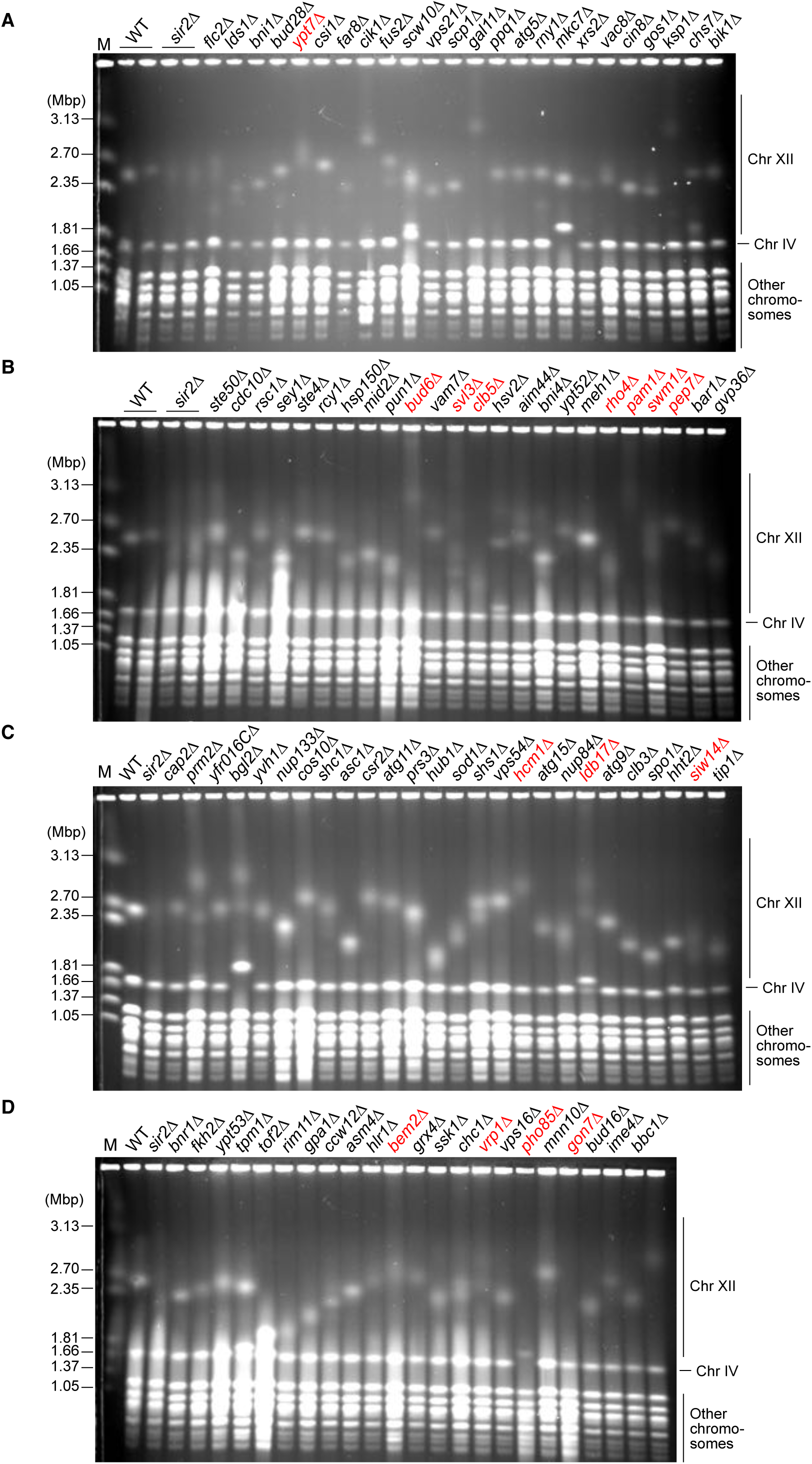
PFGE analysis of rDNA instability in the candidate strains. (A–D) The indicated mutant strains were patched from a yeast deletion library in which ∼4,800 non-essential genes are individually disrupted in the BY strain background. Cells were grown from these patches and genomic DNA was prepared and separated by PFGE. Genomic DNA was also prepared from two independent clones of WT and the *sir2*Δ mutant. DNA was stained with ethidium bromide. M indicates *H. wingei* chromosomal DNA markers. The mutants that were subjected to further analysis are indicated in red.

**Supplemental Figure 2.**
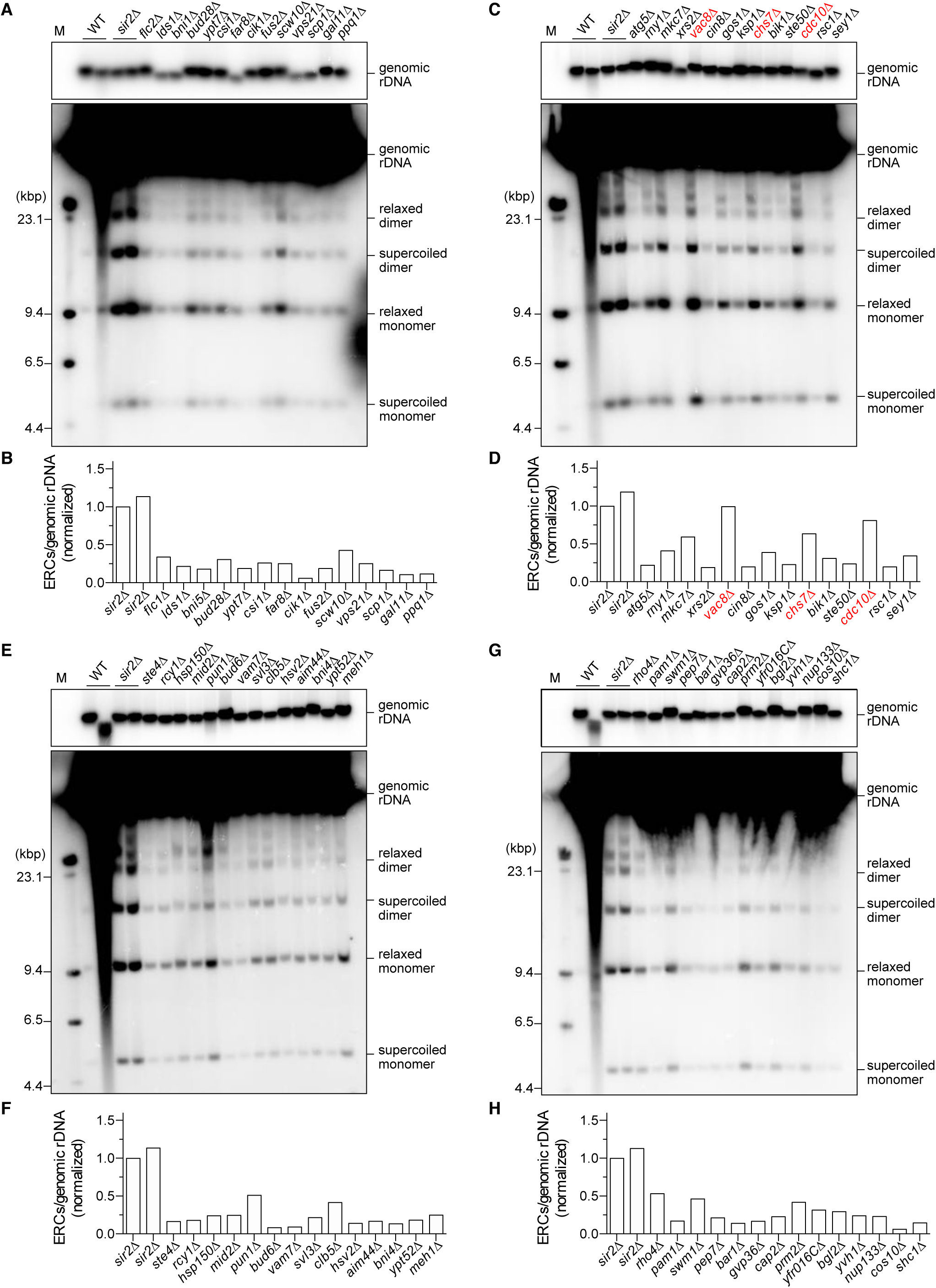
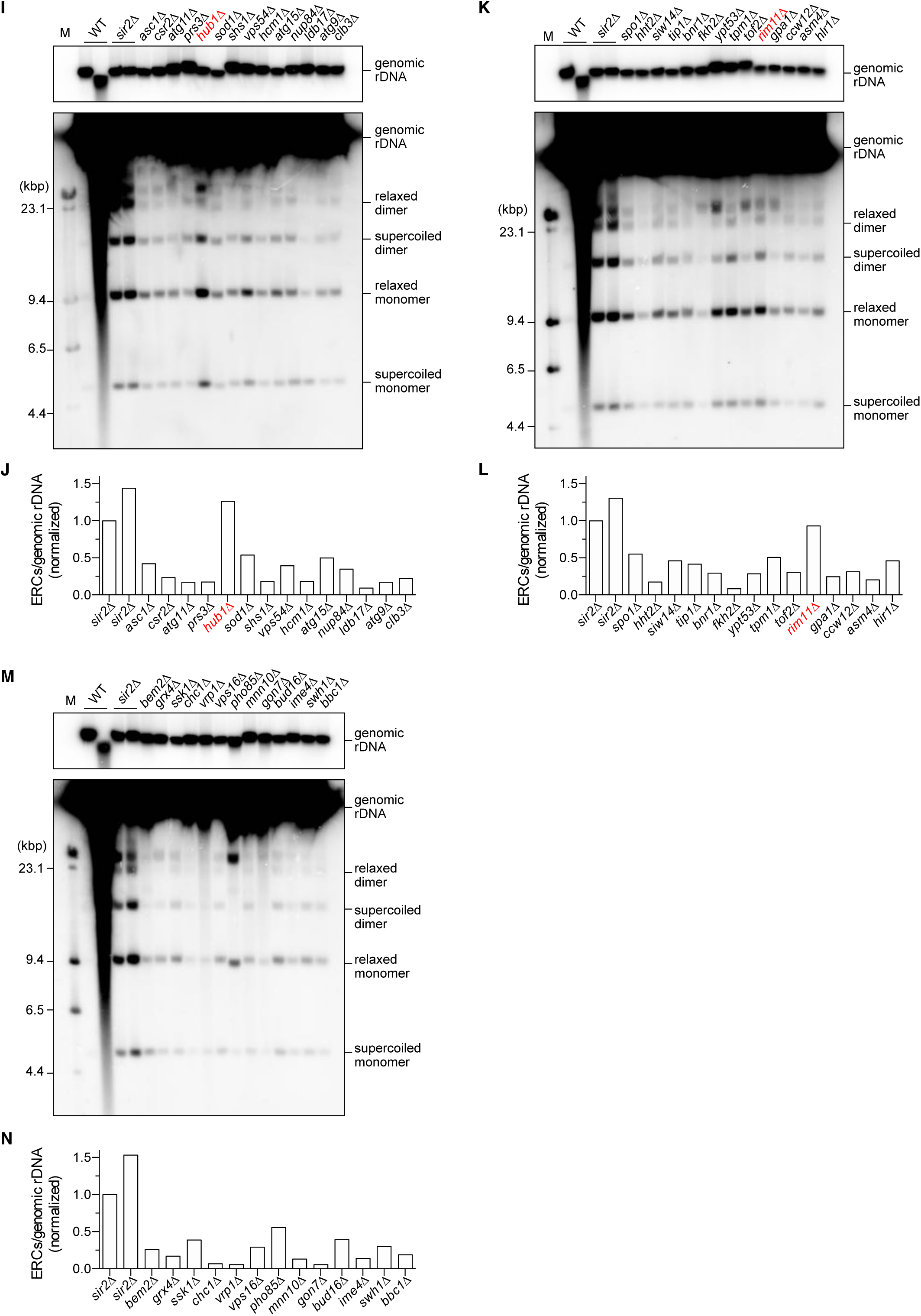
ERC analysis of rDNA instability in the candidate strains. (A, C, E, G, I, K, M) Genomic DNA was prepared from the indicated strains in the yeast deletion library as described in Fig. S1. DNA was also extracted from two independent clones of WT and the *sir2*Δ mutant. DNA was separated by conventional agarose gel electrophoresis, followed by Southern blotting with the rDNA probe. Genomic rDNA, and supercoiled and relaxed forms of monomeric and dimeric ERCs are indicated. M indicates λ DNA-*Hind* III markers. (B, D, F, H, J, L, M) Levels of total monomers and dimers relative to genomic rDNA. ERC bands were not clearly detected in one WT clone due to poor sample quality and thus quantification of WT is not shown in the graph. The level of ERCs in each mutant was normalized to that of one of the *sir2*Δ samples. The mutants that were subjected to further analysis are indicated in red.

**Supplemental Figure 3.**
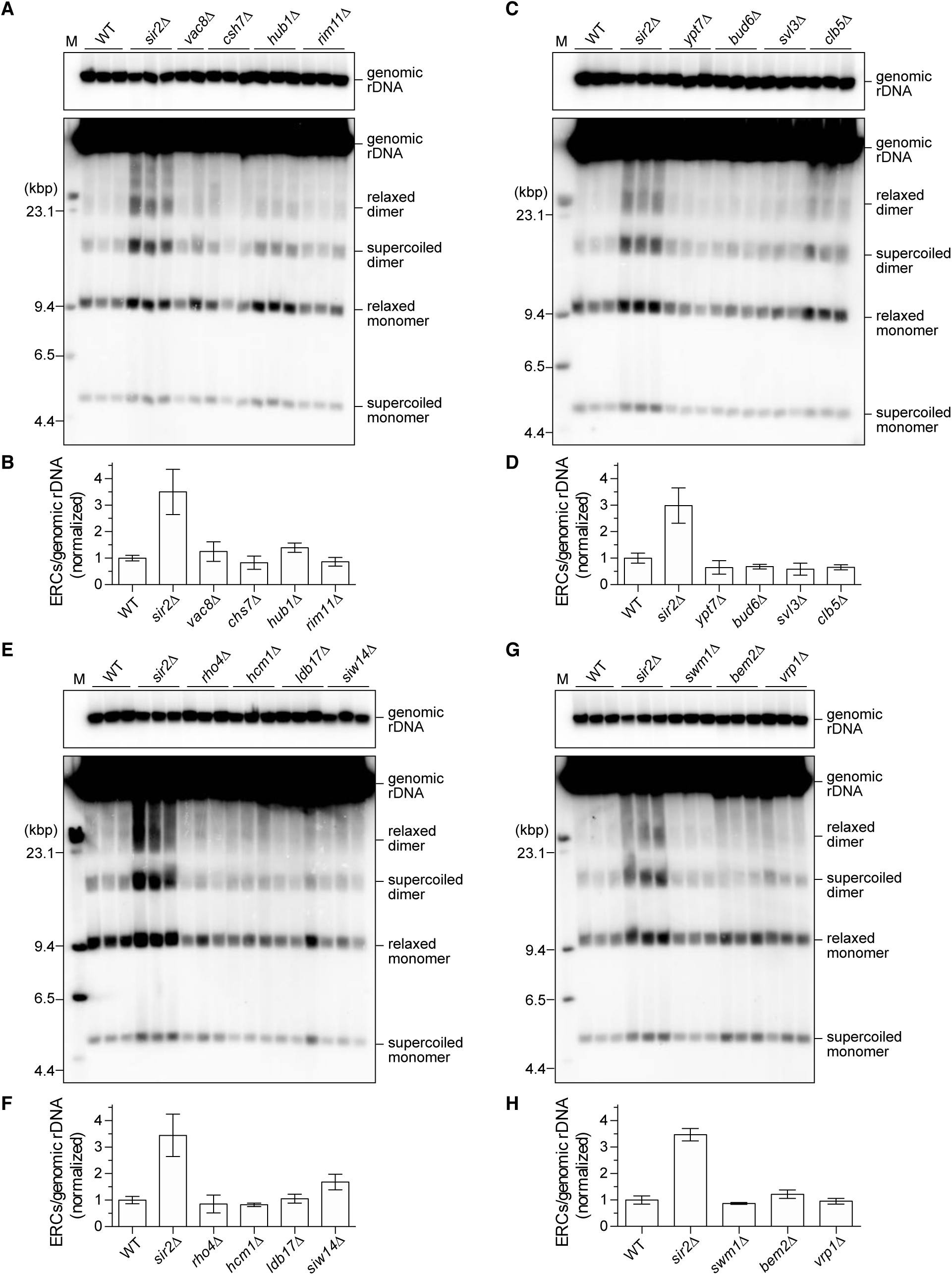
ERC analysis to examine rDNA instability in the candidate mutants in the W303 background. (A, C, E, G) The indicated genes were deleted from WT cells of the W303 background. DNA was extracted from three independent clones of WT and all mutant strains except for *vac8*Δ (two clones), and was separated by conventional agarose gel electrophoresis, followed by Southern blotting with the rDNA probe. Genomic rDNA, and supercoiled and relaxed forms of monomeric and dimeric ERCs are indicated. M indicates λ DNA-Hind III markers. (B, D, F, H) Level of total monomers and dimers relative to genomic rDNA. The level of ERCs in each mutant was normalized to the average of the WT clones (bars show mean ± SD, except for the bar of *vac8*Δ, which shows the range of two independent clones).

**Supplemental Figure 4.**
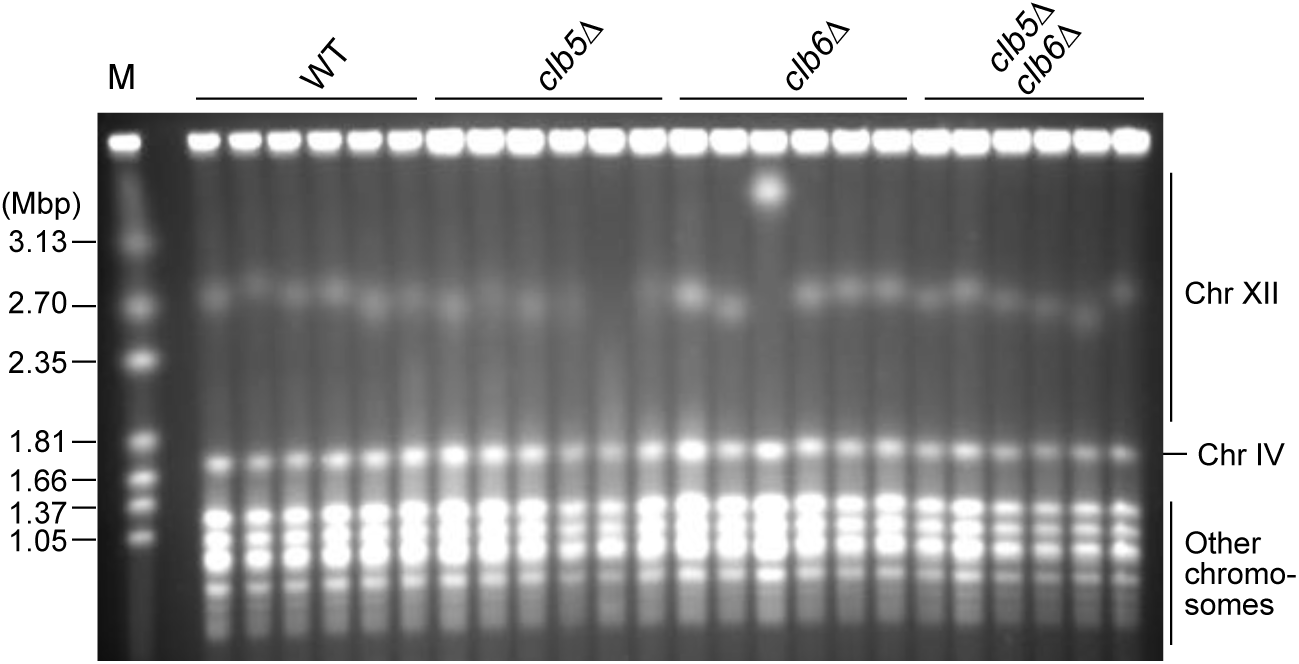
PFGE analysis of rDNA instability in the indicated strains. DNA was extracted from six independent clones of the indicated strains and separated by PFGE. DNA was stained with ethidium bromide. M indicates *H. wingei* chromosomal DNA markers.

## REFERENCES

1. Mirkin E V., Mirkin SM. 2007. Replication Fork Stalling at Natural Impediments. Microbiol Mol Biol Rev 71:13–35.

2. Branzei D, Foiani M. 2010. Maintaining genome stability at the replication fork. Nat Rev Mol Cell Biol.

3. Dewar JM, Walter JC. 2017. Mechanisms of DNA replication termination. Nat Rev Mol Cell Biol 18:507–516.

4. Liu P, Carvalho CMB, Hastings PJ, Lupski JR. 2012. Mechanisms for recurrent and complex human genomic rearrangements. Curr Opin Genet Dev 22:211–220.

5. Kobayashi T. 2011. Regulation of ribosomal RNA gene copy number and its role in modulating genome integrity and evolutionary adaptability in yeast. Cell Mol Life Sci 68:1395–1403.

6. Linskens MH, Huberman JA. 1988. Organization of replication of ribosomal DNA in Saccharomyces cerevisiae. Mol Cell Biol 8:4927–4935.

7. Kobayashi T, Hidaka M, Nishizawa M, Horiuchi T. 1992. Identification of a site required for DNA replication fork blocking activity in the rRNA gene cluster in Saccharomyces cerevisiae. MGG Mol Gen Genet 233:355–362.

8. Kobayashi T, Horiuchi T. 1996. A yeast gene product, Fob1 protein, required for both replication fork blocking and recombinational hotspot activities. Genes to Cells 1:465–474.

9. Brewer BJ, Lockshon D, Fangman WL. 1992. The arrest of replication forks in the rDNA of yeast occurs independently of transcription. Cell.

10. Kobayashi T, Horiuchi T, Tongaonkar P, Vu L, Nomura M. 2004. SIR2 regulates recombination between different rDNA repeats, but not recombination within individual rRNA genes in yeast. Cell 117:441–453.

11. Burkhalter MD, Sogo JM. 2004. rDNA enhancer affects replication initiation and mitotic recombination: Fob1 mediates nucleolytic processing independently of replication. Mol Cell 15:409–421.

12. Weitao T, Budd M, Hoopes LLM, Campbell JL. 2003. Dna2 helicase/nuclease causes replicative fork stalling and double-strand breaks in the ribosomal DNA of Saccharomyces cerevisiae. J Biol Chem 278:22513–22522.

13. Saka K, Takahashi A, Sasaki M, Kobayashi T. 2016. More than 10% of yeast genes are related to genome stability and influence cellular senescence via rDNA maintenance. Nucleic Acids Res 44:4211–4221.

14. Fragkos M, Ganier O, Coulombe P, Méchali M. 2015. DNA replication origin activation in space and time. Nat Rev Mol Cell Biol 16:360–374.

15. Donaldson AD, Raghuraman MK, Friedman KL, Cross FR, Brewer BJ, Fangman WL. 1998. CLB5-dependent activation of late replication origins in S. cerevisiae. Mol Cell 2:173–182.

16. Epstein CB, Cross FR. 1992. CLB5: A novel B cyclin from budding yeast with a role in S phase. Genes Dev 6:1695–1706.

17. Kühne C, Linder P. 1993. A new pair of B-type cyclins from Saccharomyces cerevisiae that function early in the cell cycle. EMBO J 12:3437–3447.

18. Schwob E, Nasmyth K. 1993. CLB5 and CLB6, a new pair of B cyclins involved in DNA replication in Saccharomyces cerevisiae. Genes Dev 7:1160–1175.

19. Winzeler EA, Shoemaker DD, Astromoff A, Liang H, Anderson K, Andre B, Bangham R, Benito R, Boeke JD, Bussey H, Chu AM, Connelly C, Davis K, Dietrich F, Dow SW, El Bakkoury M, Foury F, Friend SH, Gentalen E, Giaever G, Hegemann JH, Jones T, Laub M, Liao H, Liebundguth N, Lockhart DJ, Lucau-Danila A, Lussier M, M’Rabet N, Menard P, Mittmann M, Pai C, Rebischung C, Revuelta JL, Riles L, Roberts CJ, Ross-MacDonald P, Scherens B, Snyder M, Sookhai-Mahadeo S, Storms RK, Véronneau S, Voet M, Volckaert G, Ward TR, Wysocki R, Yen GS, Yu K, Zimmermann K, Philippsen P, Johnston M, Davis RW. 1999. Functional characterization of the S. cerevisiae genome by gene deletion and parallel analysis. Science (80-).

20. Wach A, Brachat A, Pöhlmann R, Philippsen P. 1994. New heterologous modules for classical or PCR-based gene disruptions in Saccharomyces cerevisiae. Yeast.

21. Giaever G, Chu AM, Ni L, Connelly C, Riles L, Véronneau S, Dow S, Lucau-Danila A, Anderson K, André B, Arkin AP, Astromoff A, El Bakkoury M, Bangham R, Benito R, Brachat S, Campanaro S, Curtiss M, Davis K, Deutschbauer A, Entian KD, Flaherty P, Foury F, Garfinkel DJ, Gerstein M, Gotte D, Güldener U, Hegemann JH, Hempel S, Herman Z, Jaramillo DF, Kelly DE, Kelly SL, Kötter P, LaBonte D, Lamb DC, Lan N, Liang H, Liao H, Liu L, Luo C, Lussier M, Mao R, Menard P, Ooi SL, Revuelta JL, Roberts CJ, Rose M, Ross-Macdonald P, Scherens B, Schimmack G, Shafer B, Shoemaker DD, Sookhai-Mahadeo S, Storms RK, Strathern JN, Valle G, Voet M, Volckaert G, Wang CY, Ward TR, Wilhelmy J, Winzeler EA, Yang Y, Yen G, Youngman E, Yu K, Bussey H, Boeke JD, Snyder M, Philippsen P, Davis RW, Johnston M. 2002. Functional profiling of the Saccharomyces cerevisiae genome. Nature.

22. Sasaki M, Kobayashi T. 2017. Ctf4 Prevents Genome Rearrangements by Suppressing DNA Double-Strand Break Formation and Its End Resection at Arrested Replication Forks. Mol Cell 66:533-545.e5.

23. Ide S, Kobayashi T. 2010. Analysis of DNA replication in Saccharomyces cerevisiae by two-dimensional and pulsed-field gel electrophoresis. Curr Protoc Cell Biol 1–12.

24. Ide S, Saka K, Kobayashi T. 2013. Rtt109 Prevents Hyper-Amplification of Ribosomal RNA Genes through Histone Modification in Budding Yeast. PLoS Genet 9.

25. Horigome C, Unozawa E, Ooki T, Kobayashi T. 2019. Ribosomal RNA gene repeats associate with the nuclear pore complex for maintenance after DNA damage. PLoS Genet 15:1–21.

26. Hosoyamada S, Sasaki M, Kobayashi T. 2019. The CCR4-NOT Complex Maintains Stability and Transcription of rRNA Genes by Repressing Antisense Transcripts. Mol Cell Biol 40:1–15.

27. Kobayashi T, Sasaki M. 2017. Ribosomal DNA stability is supported by many ‘buffer genes’-introduction to the Yeast rDNA Stability Database. FEMS Yeast Res 17:1–8.

28. Kaeberlein M, Mcvey M, Guarente L. 1999. Saccharomyces cerevisiae by two different mechanisms promote longevity in Saccharomyces cerevisiae by two different mechanisms. Genes Dev 13:2570–2580.

29. Sinclair DA, Guarente L. 1997. Extrachromosomal rDNA circles - A cause of aging in yeast. Cell 91:1033–1042.

30. Michaillat L, Mayer A. 2013. Identification of Genes Affecting Vacuole Membrane Fragmentation in Saccharomyces cerevisiae. PLoS One.

31. Friedman KL, Brewer BJ. 1995. Analysis of Replication Intermediates by Two-Dimensional Agarose Gel Electrophoresis. Methods Enzymol 262:613–627.

32. Sasaki M, Lange J, Keeney S. 2010. Genome destabilization by homologous recombination in the germ line. Nat Rev Mol Cell Biol 11:182–195.

33. Ganley ARD, Ide S, Saka K, Kobayashi T. 2009. The Effect of Replication Initiation on Gene Amplification in the rDNA and Its Relationship to Aging. Mol Cell 35:683–693.

34. Kobayashi T, Heck DJ, Nomura M, Horiuchi T. 1998. Expansion and contraction of ribosomal DNA repeats in Saccharomyces cerevisiae: Requirement of replication fork blocking (Fob1) protein and the role of RNA polymerase I. Genes Dev 12:3821–3830.

35. Brewer BJ, Fangman WL. 1988. A replication fork barrier at the 3′ end of yeast ribosomal RNA genes. Cell 55:637–643.

36. Pasero P, Bensimon A, Schwob E, Pasero P, Bensimon A, Schwob E. 2002. Single-molecule analysis reveals clustering and epigenetic regulation of replication origins at the yeast rDNA locus service Single-molecule analysis reveals clustering and epigenetic regulation of replication origins at the yeast rDNA locus 2479–2484.

37. Saffer LD, Miller OL. 1986. Electron microscopic study of Saccharomyces cerevisiae rDNA chromatin replication. Mol Cell Biol 6:1148–1157.

38. Walmsley RM, Johnston LH, Williamson DH, Oliver SG. 1984. Replicon size of yeast ribosomal DNA. MGG Mol Gen Genet.

39. Lambert S, Watson A, Sheedy DM, Martin B, Carr AM. 2005. Gross chromosomal rearrangements and elevated recombination at an inducible site-specific replication fork barrier. Cell 121:689–702.

40. Mizuno K, Lambert S, Baldacci G, Murray JM, Carr AM. 2009. Nearby inverted repeats fuse to generate acentric and dicentric palindromic chromosomes by a replication template exchange mechanism. Genes Dev 23:2876–2886.

41. Kobayashi T. 2014. Ribosomal RNA gene repeats, their stability and cellular senescence. Proc Japan Acad Ser B Phys Biol Sci 90:119–129.

42. Kobayashi T, Ganley ARD. 2005. Molecular biology: Recombination regulation by transcription-induced cohesin dissociation in rDNA repeats. Science (80-) 309:1581–1584.

43. Letessier A, Millot GA, Koundrioukoff S, Lachagès AM, Vogt N, Hansen RS, Malfoy B, Brison O, Debatisse M. 2011. Cell-type-specific replication initiation programs set fragility of the FRA3B fragile site. Nature 470:120–124.

